# Partitioning convergent modules separates phylogenetic signal from ecological information in mosaic fossils

**DOI:** 10.64898/2026.08.27.747538

**Authors:** Wei Yuan, Xuan Jing, Zi-Qiang Xu, Huateng Huang, Yanli Yue, Dong Ren, Li-Bin Ma, Jun-Jie Gu

## Abstract

Mosaic evolution assembles organisms from ancestral and derived parts, and traits shaped by convergent selection can mislead phylogenetic reconstruction while recording ecology. Disentangling these two signals within the same anatomy is a challenge for placing fossils and reconstructing evolution in deep time. We present a character-partitioning framework that fixes a molecular backbone of living species, projects fossils onto it with partitioned morphological matrices, and quantifies each anatomical module’s contribution to phylogenetic placement and ecological prediction. In mid-Cretaceous Myanmar amber crickets, which combine a cricket-like body with mole-cricket-like digging forelegs, the foreleg module drove most phylogenetic distortion and carried most ecological information: removing it restored a convergent living control species to its family and collapsed habitat-prediction accuracy from 77.8% to below the 44.4% baseline. Partitioning convergent modules thus separates phylogenetic signal from ecological information, turning mosaicism from a confound of fossil interpretation into a quantitative record of history and niche.

## INTRODUCTION

Convergent evolution shapes distant lineages under shared selective pressures (*1–4*). Because phenotypic traits evolve at heterogeneous rates, fossil phylogenetic inference is confounded by uncoupled evolutionary tempos that produce mosaic organisms and by subterranean structural adaptations that generate morphological homoplasy mimicking shared ancestry (*5–11*).

Transitioning to a fossorial lifestyle requires morphological remodeling of the appendicular skeleton to overcome soil density (*12, 13*), and this biomechanical demand drives the specialization of digging systems at the cost of functional trade-offs (*14*). Biomechanical many-to-one mapping allows lineages to achieve equivalent digging performance through alternative structural solutions (*15–17*). Because trait modules evolve at independent tempos, functional convergence can arise from varying combinations of trait complexes (*18–22*). Invading lineages either overcome phylogenetic constraints to achieve whole-body convergence or follow parallel trajectories that produce functionally analogous mosaic phenotypes. Quantitative tests of these dynamics in deep time remain lacking.

Mosaic fossils concentrate this conflict: a single specimen mixes conservative structures that track ancestry with derived modules that track ecology, and treating the two as equivalent evidence has destabilized fossil placements and the comparative inferences built upon them (*5–11*). Resolving such cases requires an analytical route that identifies which anatomical modules carry phylogenetic signal and which carry ecological information, because descriptive morphology alone cannot separate homology from homoplasy in divergent lineages.

Mid-Cretaceous orthopterans from Kachin amber, assigned to the family Pseudogryllotalpidae (*23*), embody this problem. These taxa possess fossorial prothoracic legs analogous to mole crickets (*23*) while retaining the remaining body structures of surface-dwelling crickets (Gryllidae) (*24–27*). Their familial status is contested: alternative treatments attribute the mosaic anatomy to taphonomic distortion or ontogenetic variation and synonymize the family within Gryllidae (*28, 29*), a dispute that makes these fossils a test case for partition-based placement.

We separate convergence-driven homoplasy from phylogenetic signal using a fixed molecular backbone and partitioned morphological matrices (*30, 31*). As a topological control, we employed the molecularly unsampled extant cricket *Mellogryllus mutus*, which combines derived fossorial forelegs with defining gryllid synapomorphies (*32*). Removing the foreleg partition tests the sensitivity of phylogenetic inference to localized anatomical convergence: it should restore *M. mutus* to its established classification and clarify the affinities of the Kachin amber fossils. Coupling this topological exclusion with morphospace projections separates shared ancestry from convergence (*33*) and tests whether subterranean niche colonization drives modular morphological innovation in deep time. We evaluate whether partitioning derived trait modules disentangles homoplastic ecological signals from the phylogenetic history of mosaic taxa, and what this decoupling implies for ancestral niche reconstruction.

## RESULTS

### Foreleg homoplasy obscures the phylogenetic position of fossorial lineages

Substitution saturation analysis based on the TN93 model indicated saturation at the third codon position (Figure S1). We addressed this saturation through a sensitivity framework in divergence time estimations, comparing runs with and without this position to prevent interference with deep-node branch length estimation (*34*). The topology inferred from the mitochondrial dataset served as the backbone tree because these relationships remain stable among higher Gryllidea taxa. The placement of Myrmecophilidae varies among data types (*35–38*), but this instability does not affect our conclusions: no query taxon was placed with Myrmecophilidae under any optimality criterion or matrix (Data S1). As a single query taxon, *Marchandia magnifica* mapped to the base of Gryllotalpidae across algorithms and was then constrained as a topological anchor (Data S1).

The inclusive morphological matrix grouped the eight fossil taxa as a monophyletic sister clade to extant Gryllotalpidae under Maximum Parsimony (MP), Implied Weighting (IW), and Maximum Likelihood (ML) criteria. Analyses across optimality criteria yielded divergent placements for the control species *M. mutus*. MP recovered four equally optimal placements: one at the base of Gryllotalpidae and three within or at the base of Brachytrupina, the latter consistent with its traditional classification. IW and ML recovered a single optimal placement at the base of Gryllotalpidae, with the Brachytrupina positions suboptimal (Figures 1, 2A, C, E).

**Figure 1.**
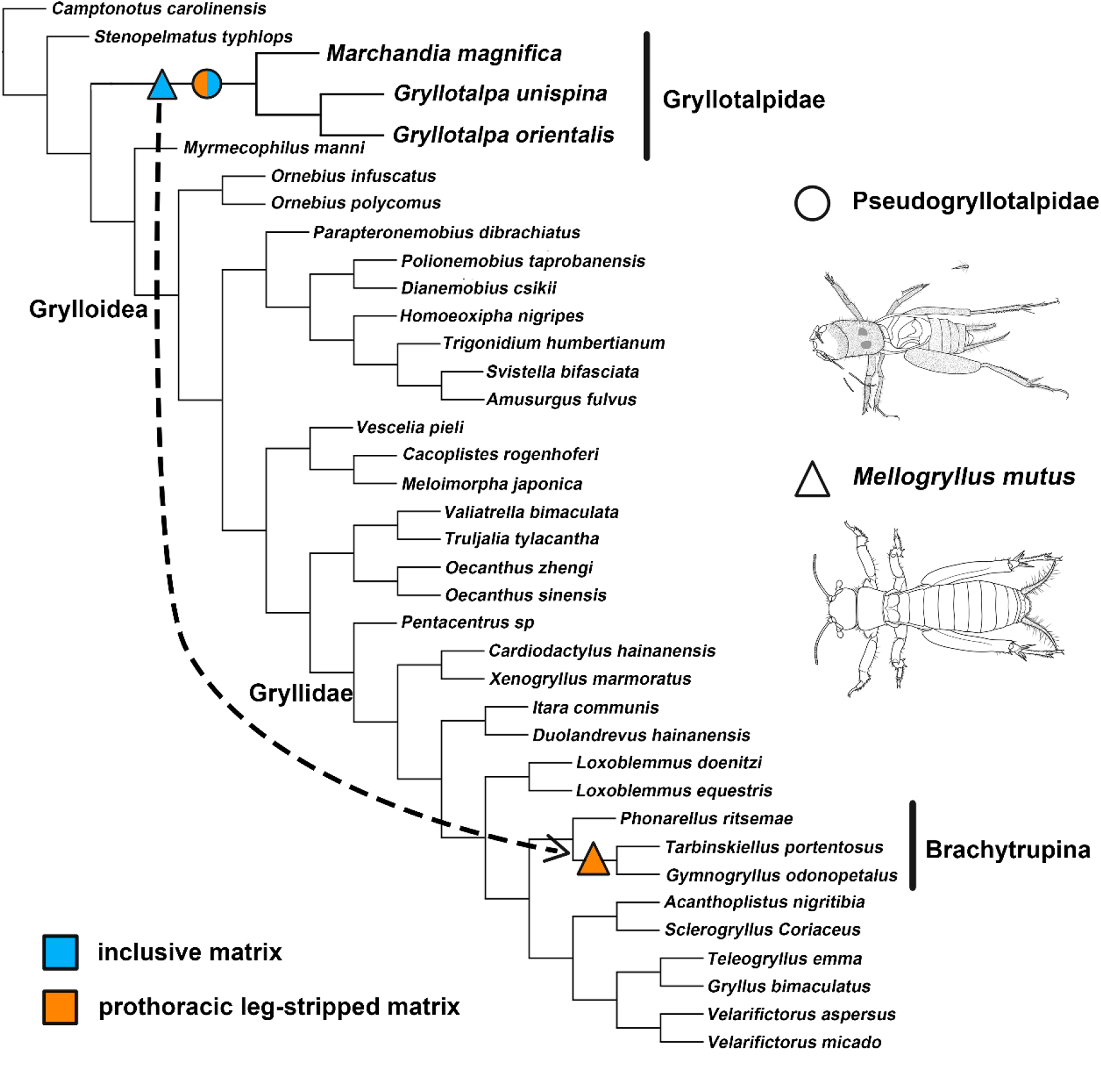
Topological dynamics of fossil and extant mosaic taxa in phylogenetic placement analyses. The cladogram represents the consensus molecular backbone tree. Symbols denote the optimal placement positions of query taxa inferred via PlaceMyFossils under different morphological matrices. The Burmese amber fossils (Pseudogryllotalpidae) are represented by a circle, and the extant control species (*M. mutus*) is represented by a triangle. Blue symbols indicate placements based on the inclusive morphological matrix (M_full), while orange symbols indicate placements based on the prothoracic leg-stripped matrix (M_stripped). The bipartite (blue/orange) circle demonstrates topological stability: the placement of Pseudogryllotalpidae remains stable at the base of Gryllotalpidae with or without the fossorial foreleg characters. The arrow connecting the blue triangle to the orange triangle illustrates a topological shift: removing the convergently evolved prothoracic leg characters resolves the morphological attraction, relocating *M. mutus* from the base of Gryllotalpidae to its expected position within the subtribe Brachytrupina.

Excluding the prothoracic leg partition induced topological divergence across optimality criteria (Data S1). MP and IW placed seven of the eight fossil species at the base of Gryllotalpidae, with several fossils also occupying alternative optimal positions within Grylloidea under MP (Figure 2B, D); the remaining species, *Chunxiania fania*, was recovered as sister to the Gryllidae crown. ML recovered only *Tresdigitus rectanguli* at the base of Gryllotalpidae, distributing the remaining seven fossil species across Mogoplistidae, Pentacentrinae, and Brachytrupina (Figure 2F). Decoupling the prothoracic leg signal leaves residual phylogenetic information sufficient to recover the Gryllotalpidae affinity of most fossils under parsimony criteria, but insufficient under likelihood criteria. The same matrix reduction separated *M. mutus* from the Gryllotalpidae clade and restored it to Brachytrupina under all three criteria, resulting in a topology congruent with traditional classification (Figures 2B, D, F; Figure 1).

**Figure 2.**
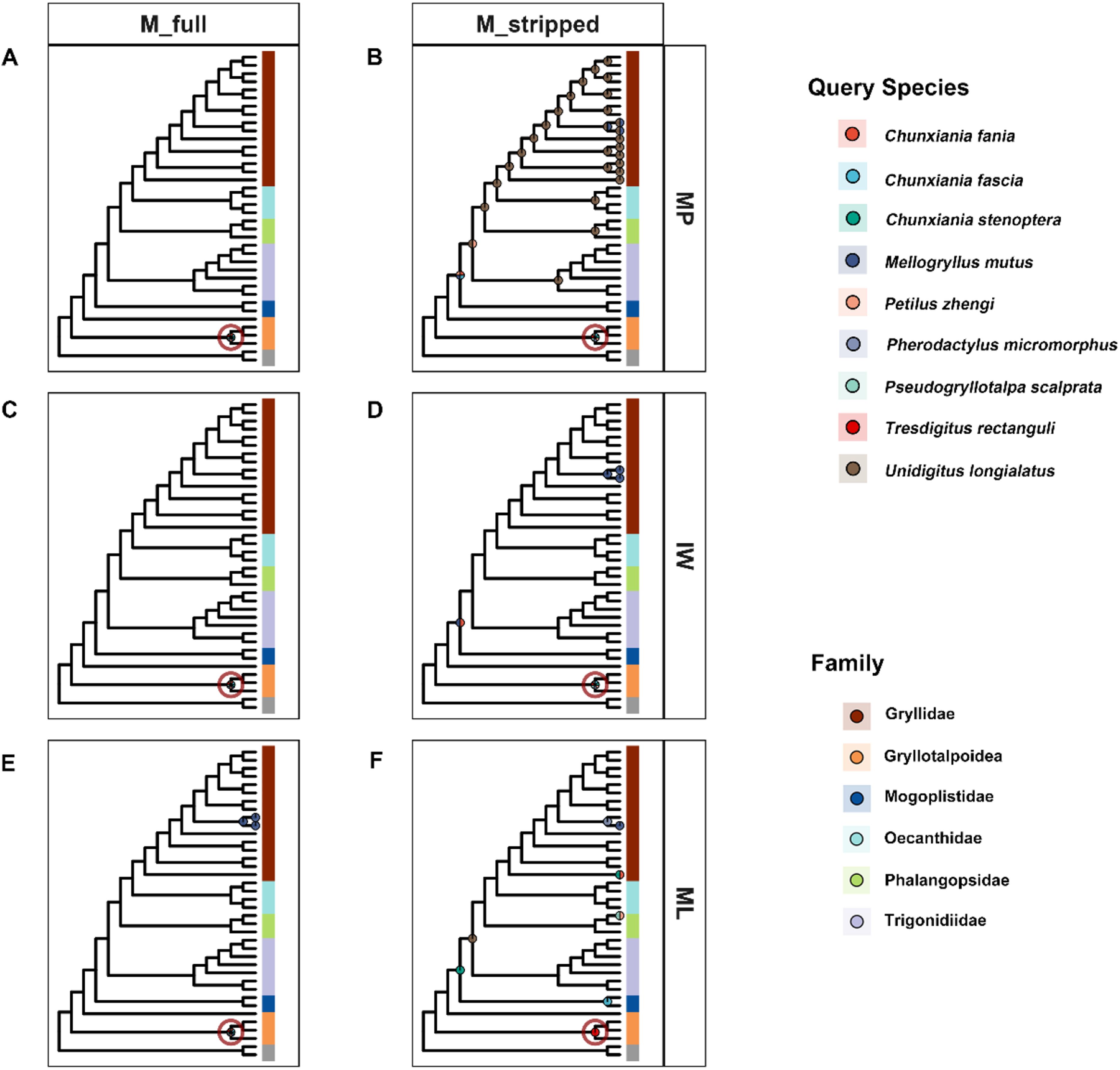
Topological mapping of optimal placements for fossil Grylloidea taxa across analytical parameters. (A, C, E) Topological placements derived from the inclusive matrix (M_full) under Maximum Parsimony (MP), Implied Weighting (IW), and Maximum Likelihood (ML) criteria, respectively. (B, D, F) Alternative optimal topologies generated using the restricted matrix excluding prothoracic leg characters (M_stripped) under MP, IW, and ML algorithms. Mapped pie charts indicate the position and relative frequency of equivalent optimal placements for each query fossil species at specific nodes. Colors correspond to the distinct query species. Solid circles represent nodes where only a single species recovered its optimal placement. The dark red open circle demarcates the reference node for the extant clade.

### Asymmetric phylogenetic signal across anatomical partitions

Evaluating the phylogenetic signal of independent morphological partitions quantified topological constraint sources. The inclusive dataset (complete) yielded the lowest Error in Placement (EP = 1.47 to 2.11). The head (15 characters) and prothoracic leg (23 characters) partitions showed the lowest errors (EP = 5.66 to 6.00 and 6.13 to 6.55, respectively) combined with high optimality score differences and low inter-taxa variance (Figure S2A, C, D), indicating that these anterior partitions dictate the placement topology. The pronotum partition (10 characters) showed intermediate errors (EP = 7.21). Errors in the metathoracic leg and abdomen partitions varied widely across query species (EP = 7.24 to 10.89 and 5.68 to 10.11), whereas the mesothoracic leg (4 characters) and wings (9 characters) partitions exceeded EP = 10 (10.11 and 10.09 to 11.59), indicating an absence of informative topological signal. In-Subtree Error (ISE) evaluations revealed topological instability within the molecular backbone concentrated at deep branches of Gryllidae, driven by conflicting signals from the wing (ISE up to 1.00) and mesothoracic leg (up to 0.93) partitions (Data S1).

### Jurassic origins of Pseudogryllotalpidae

Fossilized Birth-Death (FBD) estimations recovered the Burmese amber fossils (Pseudogryllotalpidae) as a monophyletic sister group to extant Gryllotalpidae (Figure 4A, Data S2).

Posterior probability (PP) for this relationship is sensitive to the placement of *M. mutus*, ranging from 73.48% to 75.03% in the unconstrained analyses (Runs B to D) to more than 99% when *M. mutus* is excluded (Run A, PP = 99.97%) or constrained within Gryllidae (Run E, PP = 99.73%). In the unconstrained runs, *M. mutus* was recovered as sister to the fossil plus Gryllotalpidae clade rather than within Gryllidae (Figure 3A), visualizing the morphological attraction that the constrained Run E removes. Divergence between Pseudogryllotalpidae and extant Gryllotalpidae dates to 161.34 to 198.75 Ma (95% HPD), with the fossil crown group originating between 141.52 and 184.93 Ma. The later divergence of the Brachytrupina clade containing *M. mutus* (64.08 to 110.29 Ma, Run E) demonstrates that their fossorial adaptations represent temporally and phylogenetically independent events (Figure 4A).

**Figure 3.**
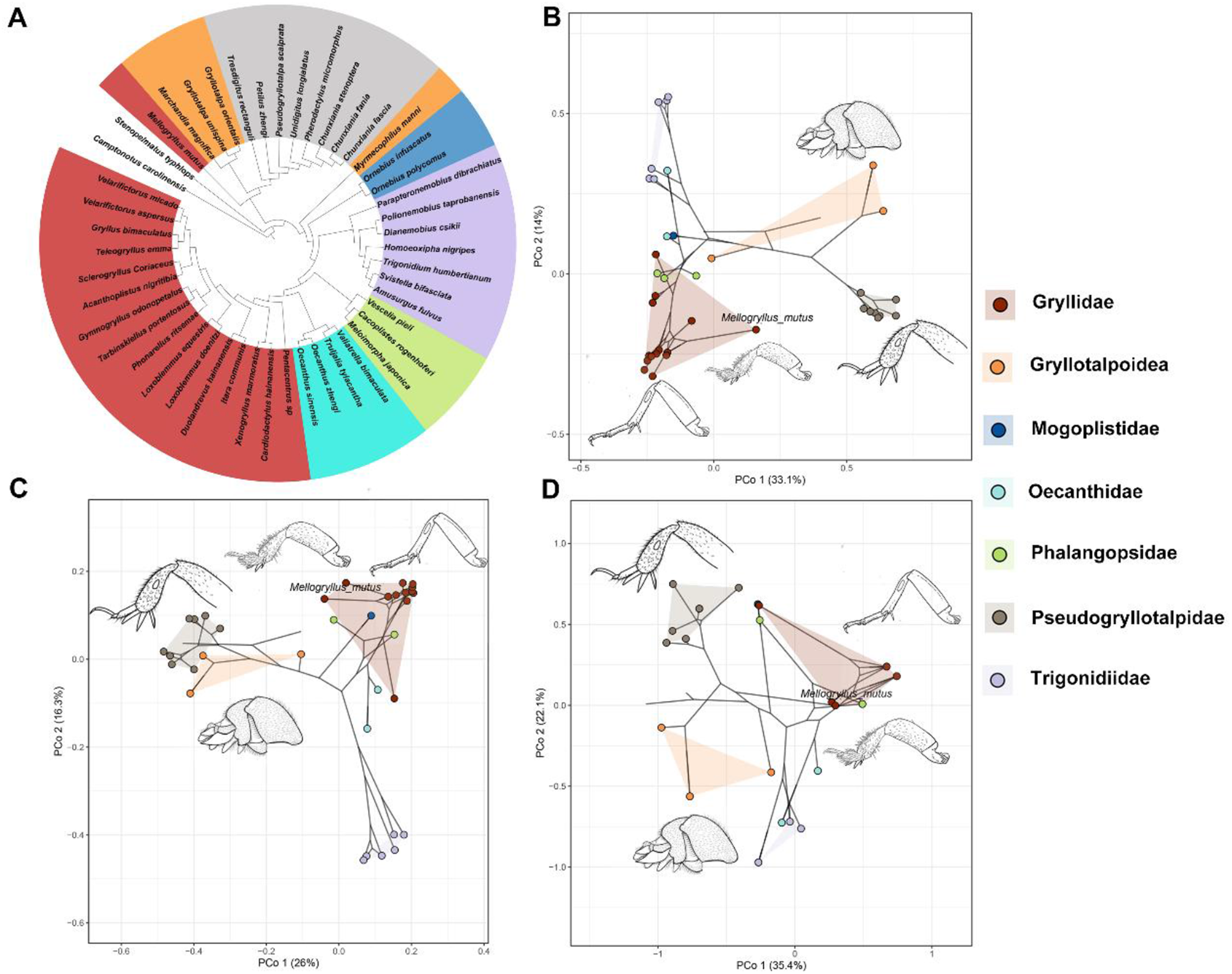
Phylogenetic framework and phylomorphospace projections of the morphological attraction driven by ecomorphological convergence. (A) Consensus cladogram derived from the Fossilized Birth-Death (FBD) analysis (corresponding to Run C). Branch lengths are omitted to emphasize topological relationships. Major family-level clades are distinguished by color. (B) Phylomorphospace constructed from the inclusive morphological matrix (M_full), mapping the phylogeny onto the first two principal coordinates (PCo1 and PCo2). (C) Phylomorphospace constructed from the prothoracic leg-stripped matrix (M_stripped). (D) Phylomorphospace constructed from the combined metathoracic leg and abdomen partition. Convex hulls delineate the morphological boundaries of the respective taxonomic groups.

**Figure 4.**
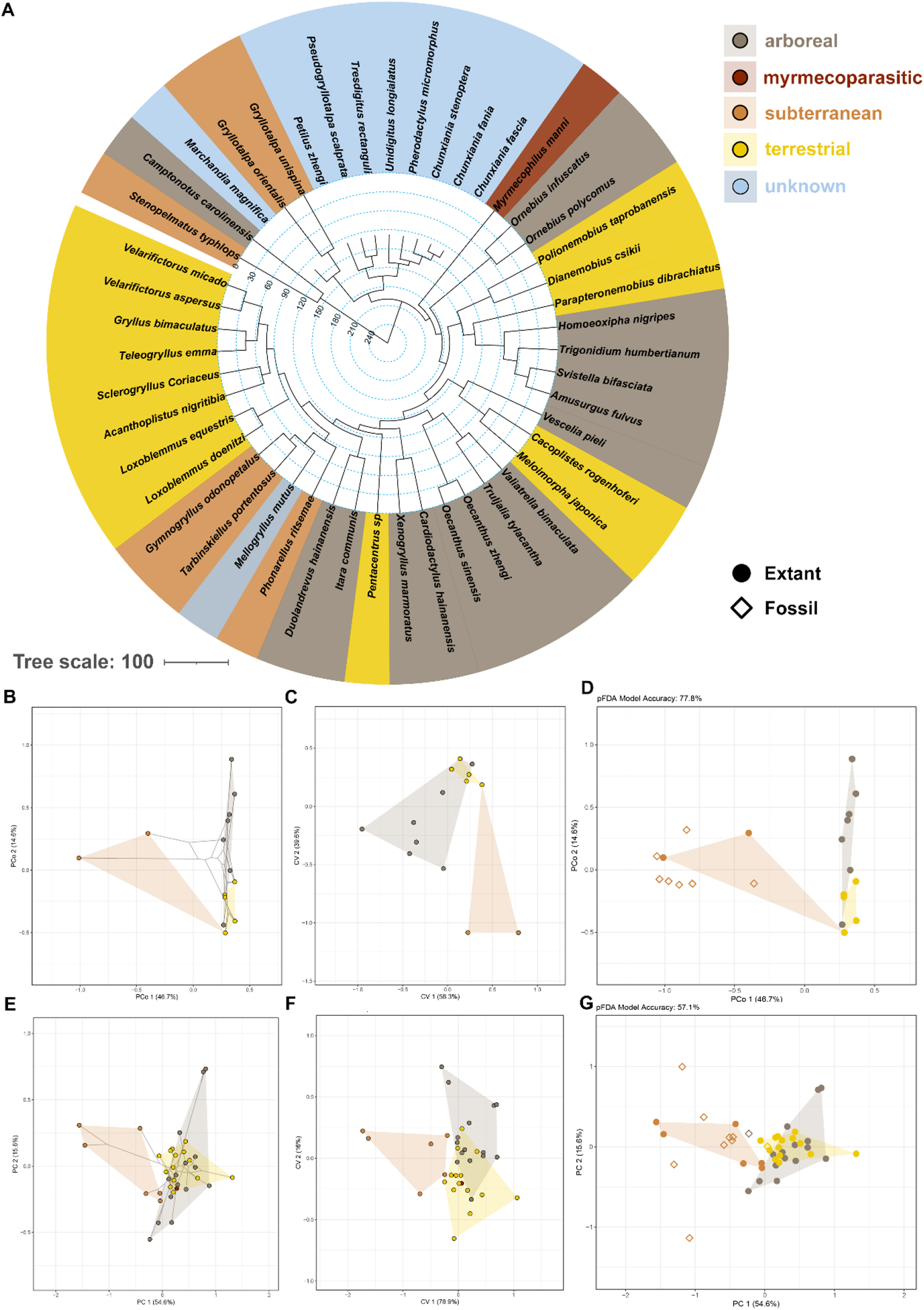
Ecomorphological dynamics and niche predictions driven by the prothoracic leg module. (A) Time-calibrated phylogenetic framework (derived from FBD Run E), with terminal taxa color-coded by microhabitat preference. (B–D) Morphological disparity and ecological discrimination based on discrete characters of the prothoracic leg partition, illustrating the (B) phylomorphospace, (C) Between-Group PCA (bgPCA), and (D) phylogenetic Flexible Discriminant Analysis (*p*FDA). Equivalent analyses (E) phylomorphospace, (F) bgPCA, and (G) *p*FDA, based on continuous linear measurements of the prothoracic leg.

### The prothoracic leg partition is the primary predictor of subterranean ecology

Principal Coordinates Analysis (PCoA) of the inclusive matrix separates fossorial lineages along the positive semi-axis of PCo1 (33.1%), distinguishing them from surface-dwelling Gryllidae and arboreal Trigonidiidae (Figure 3B). The extant control species *M. mutus* plotted at the Gryllidae margin toward Gryllotalpoidea. Excluding the prothoracic leg partition reorganizes this morphospace: fossil taxa shift toward the Gryllotalpoidea cluster on the negative semi-axis, and *M. mutus* relocates from the Gryllidae margin into the Gryllidae cluster (Figure 3C).

Principal Component Analysis (PCA) based on continuous linear measurements of the prothoracic leg partition separated subterranean taxa from arboreal and terrestrial lineages (Figure 4E–G). Phylogenetic MANOVA confirmed a microhabitat effect on prothoracic leg morphology (*R^2^* = 0.173, *F* = 2.233, *P* = 0.014; table S1). Predictive ecological modeling (pFDA) demonstrated the diagnostic value of the prothoracic leg partition. Discrete character models incorporating this partition achieved a 77.8% leave-one-out cross-validation accuracy (28 of 36 taxa; confusion matrices in table S2), predicting the eight Burmese amber fossils, *M. magnifica*, and *M. mutus* as subterranean taxa (table S1; Figure S3). Sequentially excluding the prothoracic leg partition collapsed prediction accuracy below the majority-class baseline (44.4%, 16 of 36 taxa): the Head + Pronotum model reached 38.9%, the Pronotum-only model 22.2%, and the Head-only model 16.7% (Figure 5). Under the Head-only model, the fossil taxa and *M. mutus* were still assigned to the subterranean group; given a cross-validation accuracy of 16.7%, these uniform assignments carry no ecological information (table S2; Figure S3). The Pronotum-only partition retained a group-level association with microhabitat (*R^2^* = 0.284, *P* = 0.003), whereas the Head-only partition did not reach statistical significance (*R^2^* = 0.145, *P* = 0.106; table S1).

**Figure 5.**
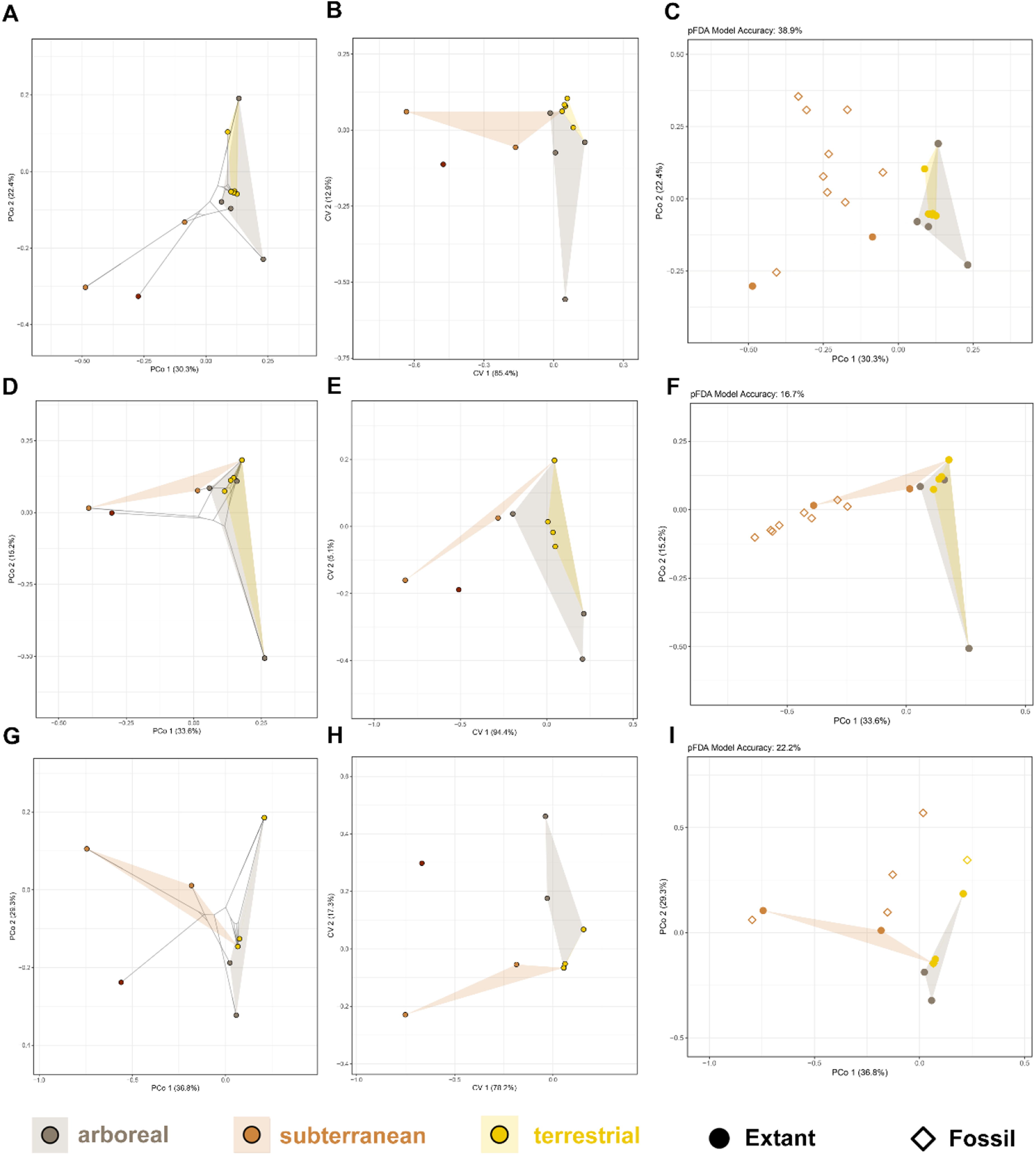
Ecomorphological dynamics and niche predictions driven by the exclusion of the prothoracic leg module. (A–C) Morphological disparity and ecological discrimination based on the combined head and pronotum partition, illustrating the (A) phylomorphospace, (B) Between-Group PCA (bgPCA), and (C) *p*FDA. (D–F) Equivalent analyses, (D) phylomorphospace, (E) bgPCA, and (F) *p*FDA, based on the head partition. (G–I) Equivalent analyses for the pronotum partition. Excluding the fossorial characters causes ecological groups to overlap.

The posterior body carries the opposing signal. The combined metathoracic leg and abdomen partition (17 characters) showed the strongest phylogenetic signal among all partitions (*K*_mult_ = 2.89, *P* = 0.001; table S1) and no significant microhabitat association (phylogenetic MANOVA: *R^2^*= 0.090, *F* = 1.084, *P* = 0.385). In the corresponding morphospace, the amber fossils form a separate cluster outside all ecological convex hulls (Figure S5I), whereas the true mole cricket *M. magnifica* falls within the subterranean hull. The pFDA model for this partition (LOOCV accuracy = 55.6%, above baseline) assigned all eight fossils and *M. mutus* to the terrestrial class (table S2). The posterior anatomy of the fossils is incompatible with the obligate subterranean plan, in direct contrast to their excavatory forelegs.

## DISCUSSION

### Mosaic assembly of subterranean adaptations

The divergence between Pseudogryllotalpidae and the extant Gryllotalpidae crown group, dated to 161.34 to 198.75 Ma (95% HPD; Data S2) via the Fossilized Birth-Death model, precedes the mid-Cretaceous amber record by at least 60 Myr. The geographic setting of this divergence cannot be inferred from present evidence. By the time these fossils were entombed in amber, however, the West Burma Block formed part of a Trans-Tethyan island arc and stood isolated from continental landmasses (*39–41*). If the lineage had been established on this block, this isolation could have provided new niches for the anterior modifications recorded in the fossils. This scenario remains speculative pending further fossil evidence.

Extant fossorial crickets exhibit diverse excavation apparatuses, ranging from the subconical heads and parallel dactylar processes of mole crickets (Figure S4A) to the widened tarsi and enlarged pulvilli of specific Gryllidae (Figure S4B, E). Cretaceous Pseudogryllotalpidae demonstrate an independent evolutionary trajectory characterized by paddle-like tibiae and apical spurs (Figure S4D). This anterior specialization evolved without corresponding modifications to the other body structures, a modular configuration corroborated by phylogenetic signal analysis. The prothoracic leg module exhibited the highest topological sensitivity, whereas other partitions maintained morphological conservatism (Figure S2). This disentangled evolution parallels the decoupled cranial and postcranial burrowing specializations of entoptychine gophers (*42*) and microteiid lizards (*43*), demonstrating that fossorial phenotypes rarely evolve as concerted whole-organism transformations (*13, 44–51*).

### Restoring the familial status of Pseudogryllotalpidae

The taxonomic treatment of the Kachin amber fossils remains contested (*23, 28, 29*). Our analyses reject the synonymy hypothesis. The eight fossil species form a monophyletic clade sister to extant Gryllotalpidae across MP, IW, ML, and all FBD runs (Figure 1, 3A; Data S1, S2). If these fossils were distorted or immature gryllids, matrix reduction should have reproduced the trajectory of the control *M. mutus*, which returned to Brachytrupina; instead, no fossil entered Gryllidae under parsimony criteria (Figure 1). The fossil clade diverged from Gryllotalpidae at 161.34 to 198.75 Ma, at least 63 Myr before the origin of Sclerogryllini (40.71 to 97.80 Ma). A sister-group position to Gryllotalpidae, combined with a foreleg architecture absent from both Gryllidae and Gryllotalpidae, excludes placement within either family. We therefore reinstate Pseudogryllotalpidae as a valid family and reject its synonymy with Sclerogryllini.

### Morphological decoupling refines ecological inference

The morphological modularity of anterior and posterior body regions refines ecological inference for extinct taxa. The foreleg and posterior partitions assign the fossils to opposite microhabitats: the excavatory forelegs predict a subterranean niche, whereas the metathoracic leg and abdomen partition shows no microhabitat association, retains the strongest phylogenetic signal of all partitions, and places the fossils outside every ecological convex hull (Figure S5I). The true mole cricket *M. magnifica* instead falls within the subterranean hull of the same morphospace. The posterior anatomy of the fossils is incompatible with the obligate subterranean plan of Gryllotalpidae, and the mosaic combination of excavatory forelegs and conservative posterior structures is more consistent with a semi-fossorial lifestyle within shallow soil or leaf litter.

Morphospace dynamics corroborate the modular evolution of these lineages. While the inclusive morphological matrix plots *M. mutus* near subterranean taxa (Figures 3B, 4, S5), excluding the prothoracic leg partition eliminates the spatial deviation driven by convergent fossorial traits, relocating *M. mutus* toward the Gryllidae cluster (Figure 3C). The Cretaceous fossil taxa follow a contrasting trajectory, shifting toward the Gryllotalpoidea cluster upon matrix reduction. This spatial divergence separates ecological convergence from shared ancestry: the fossorial phenotype of *M. mutus* reflects foreleg convergence alone, whereas the fossil morphospace shift reflects basal phylogenetic affinity unmasked by the removal of homoplastic foreleg traits. Terrestrial taxa occupy the morphospace junction between subterranean and arboreal forms (Figures 4, S5), consistent with an intermediate ecomorph from which lineages radiate under microhabitat selection pressures.

### Predictive ecological modeling and character partitioning

Predictive ecological modeling quantifies this morphological homoplasy. Models incorporating the prothoracic leg partition assign the fossil taxa to the subterranean niche (table S2; Figures 4D, S3, S5C, F). Sequential exclusion of this module collapsed cross-validation accuracy below the majority-class baseline of 44.4% (table S1; Figure 5), indicating that ecological assignments are unreliable without the prothoracic leg signal. The pronotum retains a group-level association with microhabitat during this exclusion (*R^2^* = 0.284, *P* = 0.003), consistent with its role in anchoring the foreleg musculature (*52*). This association does not translate into predictive accuracy for individual taxa (22.2%; table S1). The uniform assignment of fossil taxa to the subterranean class under the Head-only model carries no ecological information, because the underlying accuracy of 16.7% falls far below the majority-class baseline. Lacking biomechanical coupling with the excavation apparatus, the head partition provides insufficient signal for subterranean niche detection (Figure 5D–F), demonstrating that functional integration across adjacent segments limits the independence of anatomical modules.

Shared biomechanical demands drive divergent ecomorphological profiles toward equivalent functional outcomes, segregating subterranean lineages into separate morphospace regions (Figure 3B, C) (*33, 42*). This structural convergence generates an analytical conflict (*53*): fossorial forelegs confound phylogenetic inference but constitute the primary signal for predictive ecomorphology (Figures 4, 5).

A dual-matrix framework isolates the source of this homoplasy. The inclusive morphological matrix drove the attraction of *M. mutus* to the base of Gryllotalpidae; excluding the prothoracic leg module resolved this morphological attraction and restored *M. mutus* to Brachytrupina (Figure 1). While conservative signals from posterior body partitions maintain stable fossil placements under parsimony criteria (Figures 2, S2C, D), maximum likelihood models expose the limits of this residual signal: excluding the prothoracic leg disperses seven of the eight fossil species across Mogoplistidae, Pentacentrinae, and Brachytrupina (Figure 2F). Residual homoplasy in the head partition is unlikely to account for the persistent fossil placement under parsimony criteria: among all partitions, the head showed the lowest placement error (EP = 5.66 to 6.00) and the lowest subtree invasion rates (ISE < 0.4 at deep gryllid nodes), indicating backbone-congruent signal rather than homoplastic attraction (Data S1). The partitions with the highest invasion rates (wings and mesothoracic legs, ISE up to 1.00) carry too few characters to influence fossil placement.

The phenotypes of Cretaceous Pseudogryllotalpidae document a semi-fossorial ecomorph in orthopteran evolution: anterior modules adapted to subterranean excavation, whereas posterior structures retained ancestral surface-dwelling morphologies. This localized adaptation generates deep-time morphological homoplasy that acts both as a source of phylogenetic noise and as a predictor of ecological niches. Under parsimony criteria, excluding the prothoracic leg module resolves the phylogenetic distortion driven by convergent excavation traits, whereas likelihood models expose the limits of the residual signal in the reduced matrix. Retaining this module in morphospace and discriminant analyses predicts subterranean ecologies. Systematic character partitioning retrieves obscured phylogenetic histories and identifies functionally integrated modules across deep time. Although demonstrated here in amber crickets, the framework is not taxon-bound: any mosaic fossil can be decomposed into anatomical partitions, projected onto a molecular backbone of living relatives, and scored for the dual contribution of each module to phylogenetic placement and ecological prediction. Applied to other disputed mosaic fossils, for example fossorial vertebrates and burrowing arthropods whose systematic positions remain unresolved (*42, 43*), this approach converts homoplasy from an unquantifiable source of error into a quantifiable, ecologically interpretable signal in deep-time evolutionary inference.

## MATERIALS AND METHODS

### Taxon Sampling and Fossil Materials

A dataset of 47 taxa was assembled to test between convergent adaptation and phylogenetic constraint. A phylogenetic backbone was constructed using 37 extant orthopteran species, including two outgroups from the superfamily Stenopelmatoidea (*Camptonotus carolinensis* (Gerstaecker, 1860) and *Stenopelmatus typhlops* Rehn, 1903). The ingroup comprises 35 extant species representing 7 families and 16 subfamilies within the infraorder Gryllidea. Backbone taxa were selected based on two criteria: (1) the availability of complete mitogenome data to ensure a stable backbone topology; and (2) the accessibility of morphological data (*37*).

The focal taxa comprise eight fossil species from mid-Cretaceous Burmese amber (ca. 99 Ma) (*54*). Species-level taxonomy within *Chunxiania* follows a companion taxonomic study (T.-Y. Gao, W. Yuan, and J.-J. Gu, unpublished), which revalidates *C. fascia* and describes the new species *C. stenoptera*. To evaluate whether their mosaic phenotypes represent an independent evolutionary trajectory rather than taphonomic distortion or ontogenetic variation (*27, 29, 55*), these fossils were treated as independent operational taxonomic units (OTUs) a priori. The early-instar nymph *Burmagryllotalpa longa* Wang et al., 2019 was excluded to avoid ontogenetic bias. The Lower Cretaceous mole cricket *Marchandia magnifica* (*56*) was integrated as a structural and topological anchor. The extant gryllid *Mellogryllus mutus* Cadena-Castañeda et al., 2022, which possesses convergently evolved fossorial prothoracic legs but lacks molecular data, was included as a morphological proxy to test the vulnerability of phylogenetic inference to morphological convergence. The nine amber fossils were added to the 38 extant species (37 backbone species plus *M. mutus*), yielding 47 taxa in total. Extant taxa were assigned to four microhabitat categories (arboreal, terrestrial, subterranean, and myrmecoparasitic) based on published ecological records.

### Morphological Character Acquisition and Matrix Construction

Morphological data were scored through direct examination of adult extant specimens and fossil inclusions, or from published descriptions and images when specimens were unavailable. The morphological matrix consists of 78 discrete characters partitioned into seven anatomical regions (table S3): Head (1–15), Pronotum (16–25), Prothoracic leg (26–48), Mesothoracic leg (49–52), Metathoracic leg (53–62), Wings (63–71), and Abdomen (72–78). Inapplicable characters were coded as ‘-’ and missing data as ‘?’. Two distinct character matrices were established: (1) the inclusive matrix (M_full), encompassing all 78 characters; and (2) the stripped matrix (M_stripped), excluding the entire prothoracic leg partition (23 characters). Ten linear measurements of the prothoracic leg were obtained for all taxa: profemur length (mid-section), profemur width (mid-section and distal section), protibia length, protibia width (base, mid, and distal sections), probasitarsus length, probasitarsus width, and claw length, derived from high-resolution photographs or scaled images and descriptions in the primary literature (*24–27, 57, 58*).

### Molecular Data and Substitution Saturation Analysis

Mitochondrial sequences for the outgroups and 36 ingroup species were obtained from GenBank and Jing et al. (*37*). A substitution saturation analysis was performed by plotting the observed number of transitions and transversions against genetic distances calculated under the Tamura-Nei (TN93) model (*59*). This analysis identified saturation at the third codon position, which was addressed through a sensitivity framework during divergence time estimation.

### Phylogenetic Placement Analysis

We used PlaceMyFossils (*30*) in TNT v1.6 (*60*) to project query taxa onto an extant molecular backbone. The backbone tree was inferred using IQ-TREE 2 (*61*); best-fit substitution models and partitioning were selected via ModelFinder (*62*) based on BIC scores, with nodal support assessed using 1,000 ultrafast bootstrap replicates. The fossil *M. magnifica* was first analyzed as a single query and then constrained as a topological anchor. Following this validation, nine target query species (eight Burmese amber fossils and *M. mutus*) were analyzed using a stepwise evaluation strategy:

1. Basic placement mapping onto the fixed molecular backbone using both M_full and M_stripped;
2. Independent placement under Maximum Parsimony (MP), Implied Weighting (IW), and Maximum Likelihood (ML) utilizing both matrices to assess placement stability and distinguish true phylogenetic signal from ecomorphological convergence (*30, 63, 64*). For IW, the optimal concavity constant (k) was determined independently for each query species via the automated Leave-One-Out Validation (LOOV) routine implemented within the PlaceMyFossils pipeline, evaluating a predefined parameter space ranging from k=3 to k=30 across 10 discrete intervals.
3. Character partition analysis to evaluate specific anatomical contributions.
4. Error in Placement (EP) and In-Subtree Error (ISE) calculations. EP approximates the ability of the query dataset to recover known relationships: each backbone species was pruned from the reference tree and re-placed using only the query dataset, and EP equals the mean nodal distance between the original and optimal positions across backbone species, using the maximum nodal distance when multiple optimal positions occurred (*30*). EP was calculated for the complete dataset and for each character partition separately. The ISE quantifies, for each subtree of the reference tree, the fraction of backbone species not belonging to that subtree that were optimally placed within it, flagging backbone regions where placements are unreliable (*30*). The “Use char. sampling of target species” function was activated to match the missing data profile of each respective query fossil, approximating empirical phylogenetic instability.

### Divergence Time Estimation

Divergence times were estimated using the Fossilized Birth-Death (FBD) model in MrBayes v. 3.2.7 (*65, 66*). Monophyly constraints were applied to extant major clades supported by molecular data. No hard constraints were applied to any fossil taxa, including *M. magnifica*. The molecular dataset was partitioned by codon position (best-fit models via ModelFinder). The morphological data were partitioned anatomically into a non-prothoracic leg partition (55 characters) and a prothoracic leg partition (23 characters), each modeled under the Mkv+Γ model with unlinked parameters and rate multipliers (ratepr=variable). Priors followed orthopteran-specific calibrations (*35, 67*): the clock used independent gamma rates (IGR) with exponential hyperprior (rate parameter=2, mean=0.5); speciation rate was set to exponential prior (rate parameter=10, mean=0.1); extinction rate followed a Beta (1,1) distribution. *Protogryllus grandis* (189.8-201.5 Ma) calibrated the Gryllidea node (*68*). A stepwise series of five FBD analyses (50 million generations each, sampled every 5000 generations, assessed via Tracer v1.7 (*69*)) was executed to quantify trait-driven topological bias: Run A (M_full, excluding *M. mutus*); Run B (M_full, including *M. mutus* with unpartitioned morphology); Run C (M_full, including *M. mutus* with partitioned morphology); Run D (identical to Run C but excluding saturated third codon positions); and Run E (identical to Run D but enforcing the consensus systematic position of *M. mutus* within Gryllidae). In Run E, *M. mutus* was constrained to Gryllidae (Brachytrupina). Its taxonomic placement relies on male genitalia (*32*), structures decoupled from the fossorial prothoracic legs, so constraining the phylogeny with the non-prothoracic partition isolates these traits. The topology and divergence times from Run E served as the reference phylogenetic backbone for comparative methods accounting for phylogenetic non-independence. To visualize topological distortion driven by morphological attraction, the tree from Run C was projected onto the morphospaces.

### Morphospace Analysis

All statistical analyses were conducted in R v4.4.0. To quantify morphological disparity between M_full and M_stripped matrices, Principal Coordinates Analysis (PCoA) was performed using pairwise Maximum Observable Rescaled Distance (MORD) metrics implemented in the Claddis package (*70*). The time-calibrated tree (Run C) was projected onto the M_full, M_stripped, and posterior-module morphospaces to evaluate convergent evolutionary trajectories. Taxa were classified into seven taxonomic categories: Mogoplistidae, Trigonidiidae, Oecanthidae, Gryllidae, Phalangopsidae, Gryllotalpoidea, and Burmese amber fossil taxa. Convex hulls representing these groups were generated using ggplot2 (*71*). For microhabitat-based comparisons, convex hulls were delineated by microhabitat category instead.

For continuous measurements of the prothoracic leg, *Camptonotus carolinensis* was excluded because all ten measurements were unavailable, yielding a dataset of 46 taxa × 10 variables containing 12 missing values. Missing values were imputed from the raw measurements using a random forest algorithm via the missForest package (*72*). The imputed dataset was log-transformed and converted to Log Shape Ratios (LSRs) via Mosimann transformation (*73*) to remove isometric size effects prior to Principal Component Analysis (PCA). Recognizing that subterranean adaptation imposes biomechanical constraints on the anterior functional module, our analysis focused on the head, pronotum, and prothoracic leg (*42, 52, 74, 75*). Part of the subsequent methodology follows Bignon et al. (*33*).

### Ecomorphological Modeling and Phylogenetic Signal

Phylogenetic signal was assessed for individual linear traits (Blomberg’s *K*) and multivariate shape variables (*K*_mult_) (*76, 77*). For discrete datasets, constant characters were removed, and the evolutionary conservatism of individual binary characters was evaluated using the phylogenetic *D*-statistic (*78*) via the caper package (*79*). To test microhabitat effects while accounting for phylogenetic non-independence, an RRPP-based phylogenetic MANOVA (999 iterations) was applied to the PC and PCoA scores (*80, 81*). Axis-specific morphological divergence was evaluated using univariate phylogenetic ANOVAs (*77*), and group separation was visualized via Between-Group PCA (*82*).

Ecological groups were predicted using phylogenetic Flexible Discriminant Analysis (*p*FDA) (*83*). For the discrete character partitions, *p*FDA was applied to a set of PCoA axes explaining ≥ 90% of the cumulative variance (a maximum of four axes). For the continuous measurements, the number of retained PC axes was optimized: models were fitted starting from the maximum number of axes and reduced stepwise until a non-singular fit was obtained, retaining nine PCs. In both cases, the phylogenetic signal parameter (*λ*) was optimized by grid search over [0, 1] to minimize the residual sum of squares and prevent matrix singularity and overfitting. Predictive accuracy was validated using Leave-One-Out Cross-Validation (LOOCV) and evaluated against the majority-class baseline (44.4%, 16 of 36 modeled taxa). The discrete morphological analyses and predictive modeling were executed across the following partition strategies: (1) functionally inclusive modules (Head + Pronotum + Prothoracic leg, Pronotum + Prothoracic leg, and Prothoracic leg only); (2) exclusively non-fossorial modules (Head + Pronotum, Head only, Pronotum only); and (3) a posterior module combining the metathoracic leg and abdomen partitions (17 characters). Taxa with unplaceable PCoA scores and ecological classes containing fewer than four representative taxa were excluded from modeling.

### Data and materials availability

All phylogenetic trees, morphological matrices (including M_full and M_stripped), and raw linear measurement datasets have been deposited at Zenodo and are publicly available. DOI: 10.5281/zenodo.22113815. Mitochondrial sequences used in this study are available from GenBank under accessions NC_028060.1 and NC_072277.1 and from Jing et al. (*37*).

Original R scripts used for morphospace projection, phylogenetic MANOVA, and *p*FDA have been deposited at Zenodo and are publicly available.

## Author contributions

Conceptualization, W.Y., L.-B.M., and J.-J.G.; Methodology, W.Y. and Z.-Q.X.; Investigation, W.Y., Z.-Q.X., and J.-J.G.; Formal analysis, W.Y.; Resources, L.-B.M., X.J., H.-T.H. and J.-J.G.; Visualization, W.Y.; Funding acquisition, L.-B.M and J.-J.G.; Supervision, L.-B.M., H.-T.H., and J.-J.G.; Writing - original draft, W.Y. and J.-J.G.; Writing - review & editing, L.-B.M., H.-T.H., Y.- L.Y., D.R., and J.-J.G.

## SUPPLEMENTARY INFORMATION

Table S1. Phylogenetic signal and phylogenetic MANOVA and phylANOVA statistics for each morphological partition and for continuous prothoracic leg measurements. Available at Zenodo (https://doi.org/10.5281/zenodo.22113815).

Table S2. Observed and pFDA-predicted microhabitat assignments for all modeled taxa across partition strategies. Available at Zenodo (https://doi.org/10.5281/zenodo.22113815).

Table S3. List of characters used for the phylogenetic analysis. Available at supplementary materials.

Data S1. PlaceMyFossils output files: molecular backbone tree, basic placement mappings, and independent placement analyses under MP, IW, and ML criteria. Available at Zenodo (https://doi.org/10.5281/zenodo.22113815).

Data S2. MrBayes Fossilized Birth-Death divergence-time analyses (Runs A–E) and IQ-TREE backbone inference files. Available at Zenodo (https://doi.org/10.5281/zenodo.22113815).

