## Supplementary material for "Partitioning convergent modules separates phylogenetic signal from ecological information in mosaic fossils": Table S3 and Figures S1 to S5

**This PDF file includes:**

Table S3

Figures S1 to S5

References (S1, S2)

**Table S3. List of characters used for the phylogenetic analysis**

1. Head covered with scales: 0. yes; 1. no
2. Head: 0. Hypognathous; 1. subprognathous; 2. prognathous
3. Head with long setae: 0. yes; 1. no
4. Head with punctures: 0. yes; 1. no
5. Head with sclerotized processes: 0. yes; 1. no
6. If head has sclerotized processes, orientation of processes: 0. Longitudinal; 1. Transverse
7. If head has sclerotized processes, presence of a median facial groove: 0. yes; 1. no
8. If head has sclerotized processes, head flattened: 0. yes; 1. no
9. Clypeus situated between the antennae: 0. yes; 1. no
10. Clypeus swollen: 0. yes; 1. no
11. Frontal process prominent: 0. yes; 1. no
12. Vertex with longitudinal band: 0. yes; 1. no
13. Vertex joined with pronotum: 0. yes; 1. no
14. Length of Antennal flagella: 0. Longer than body length; 1. shorter than body length
15. Shape of antennal flagellomeres: 0. uniform and cylindrical; 1. non-cylindrical or irregular
16. Width of pronotum compared to head: 0. narrower than head; 1. nearly equal; 2. wider than head
17. Pronotum raised: 0. yes; 1. no
18. Pronotum covered with scales: 0. yes; 1. no
19. Pronotum with punctures: 0. yes; 1. no
20. Pronotum disc covered with long setae: 0. yes; 1. no
21. Pronotum with circular spots: 0. yes; 1. no
22. Lateral margins of pronotum: 0. straight; 1. curved
23. Lateral margins of pronotum gradually widening: 0. yes; 1. no
24. Pronotum covering base of forewings: 0. yes; 1. no
25. Posterior margin of pronotum narrowed: 0. yes; 1. no
26. Protrochanter prominent: 0. yes; 1. no
27. Width of profemur vs. protibia: 0. nearly equal; 1. profemur wider than protibia
28. Profemur length/width ratio > 3: 0. yes; 1. no

29. Dense long setae present on outer side of protibia: 0. yes; 1. no
30. Protibia with ventral spurs: 0. yes; 1. no
31. Base of protibia swollen: 0. yes; 1. no
32. Protibia laterally widened: 0. yes; 1. no
33. Distal part of protibia wider than basal part: 0. yes; 1. no
34. Number of movable apical spurs on protibia: 0. one; 1. two; 2. three; 3. four; 4. five
35. Presence of immovable apical spurs: 0. yes; 1. no
36. Arrangement of apical spurs: 0. in a single row; 1. not single row
37. Apical spurs longer than probasitarsus: 0. yes; 1. no
38. Basitarsus of prothoracic legs wider than that of mesothoracic legs: 0. yes; 1. no
39. Shape of basitarsus: 0. cylindrical; 1. transversely compressed ventrally; 2. flattened
40. Pulvillus structure: 0. ridge-like, 1. thickened pad-like, 2. non-thickened pad-like
41. Basitarsus with: 0. setae, 1. triangular spines
42. Second tarsomere: 0. subcylindrical, 1. laterally compressed
43. Pulvillus shape of Second tarsomere: 0. cylindrical; 1. transversely compressed ventrally; 2. flattened
44. Second tarsomere with triangular spines: 0. yes; 1. no
45. Pulvillus of second tarsomere with ventral hook-like setae: 0. yes; 1. no
46. Claws inner margin serrulated: 0. yes; 1. no
47. Claws bifid: 0. yes; 1. no
48. Number of prothoracic tarsomeres: 0. three tarsomeres; 1. four tarsomeres
49. Mesotibia with spines: 0. yes; 1. no
50. Presence of ventral apical spur on mesotibia: 0. yes; 1. no
51. Number of apical spurs on mesotibia: 0. ≤2; 1. 3–4; 2. ≥5
52. Number of mesothoracic tarsomeres: 0. three tarsomeres; 1. four tarsomeres
53. Length-to-width ratio of metafemur: 0. ≤3; 1. 4–5; 2. ≥6
54. Ventral margin of metafemur: 0. straight, 1. curved
55. Subapical spurs present on inner dorsal side of metatibia: 0. yes; 1. no
56. Subapical spurs present on outer dorsal side of metatibia: 0. yes; 1. no
57. Inner dorsal spines on metatibia gradually increasing in size toward apex: 0. yes; 1. no
58. Number of metathoracic tarsomeres: 0. three tarsomeres; 1. four tarsomeres
59. Hind basitarsus cylindrical: 0. yes; 1. no
60. Ventral surface of hind basitarsus pad-like: 0. yes; 1. no
61. Dorsal serrulation present on hind basitarsus: 0. yes; 1. no
62. Dorsal spurs present on hind basitarsus: 0. yes; 1. no
63. Wings present: 0. yes; 1. no
64. Forewing with truncate apex: 0. yes; 1. no
65. Dorsal field twice as wide as abdomen: 0. yes; 1. no
66. PCuA vein with stridulatory teeth: 0. yes; 1. no
67. Orientation of lateral field veins relative to Sc vein: 0. parallel to dorsal field; 1. perpendicular to dorsal field
68. PCuA in basal field: 0. Longitudinal extension longer than transverse; 1. Transverse extension longer than longitudinal
69. Presence of mirror: 0. yes; 1. no

70. Mirror with one internal crossvein: 0. yes; 1. no
71. Female forewings coriaceous: 0. yes; 1. no
72. Abdomen cylindrical: 0. yes; 1. no
73. Cerci length: 0. shorter than abdominal segments; 1. longer than abdominal segments but shorter than body; 2. approximately equal to body length
74. Cerci with long setae: 0. yes; 1. no
75. Ovipositor absent: 0. yes; 1. no
76. Ovipositor apex shape: 0. needle-shaped; 1. rod-shaped; 2. flattened laterally and distinctly upcurved; 3. long, cone-shaped; 4. short sword-shaped
77. If ovipositor needle-shaped: 0. straight, 1. curved
78. Dorsal margin of ovipositor apex: 0. smooth; 1. dentate; 2. tuberculate

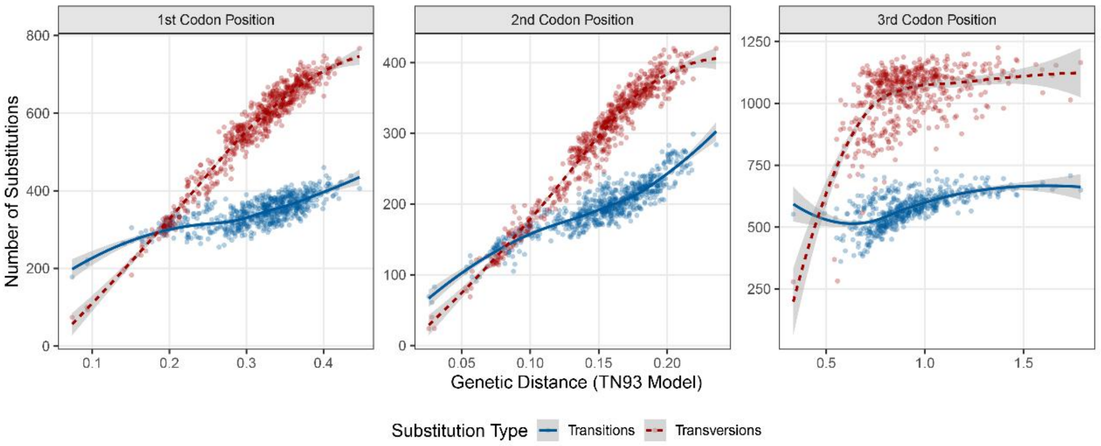


**Figure S1. Substitution saturation analysis across three codon positions.** Scatter plots display the absolute number of pairwise transitions (blue, solid line) and transversions (red, dashed line) plotted against genetic distances estimated via the Tamura-Nei 1993 (TN93) model.

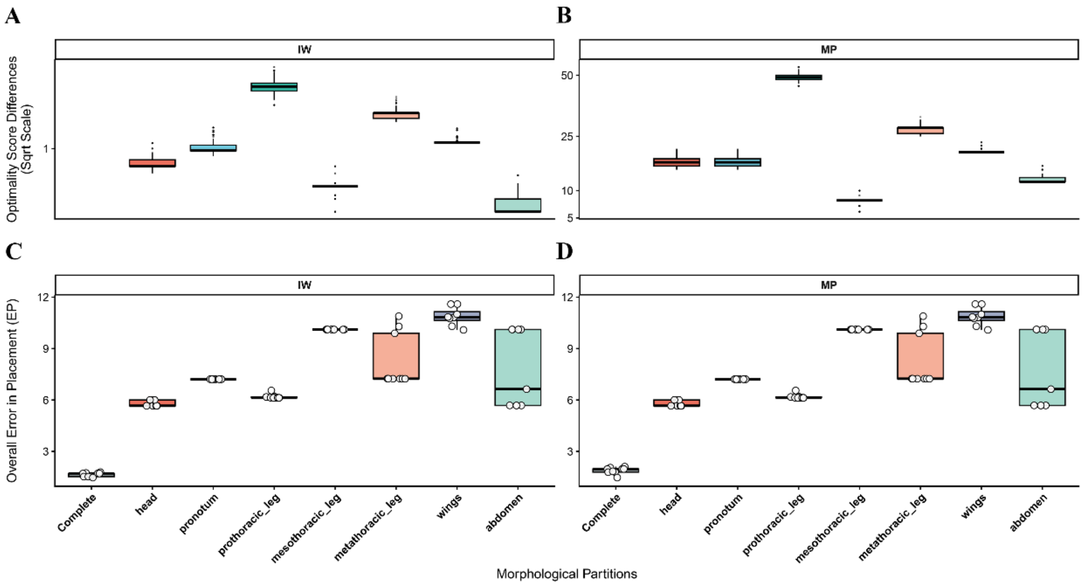


**Figure S2. Topological sensitivity and placement error across morphological partitions under**

**Maximum Parsimony (MP) and Implied Weighting (IW) criteria.** (A, B) Distribution of optimality score differences for alternative placements of query species across seven morphological partitions under (A) IW and (B) MP criteria. Under the MP criterion, scores represent the number of extra steps required for a given suboptimal placement. Under the IW criterion, scores denote the difference in total fit (fit reduction) resulting from character conflict penalization. Higher scores indicate stronger partition-specific topological constraints. (C, D) Error in Placement (EP) evaluating the topological reliability of each partition under (C) IW and (D) MP criteria. The complete dataset serves as the baseline for maximum congruence with the reference molecular backbone. Lower EP values indicate higher placement accuracy.

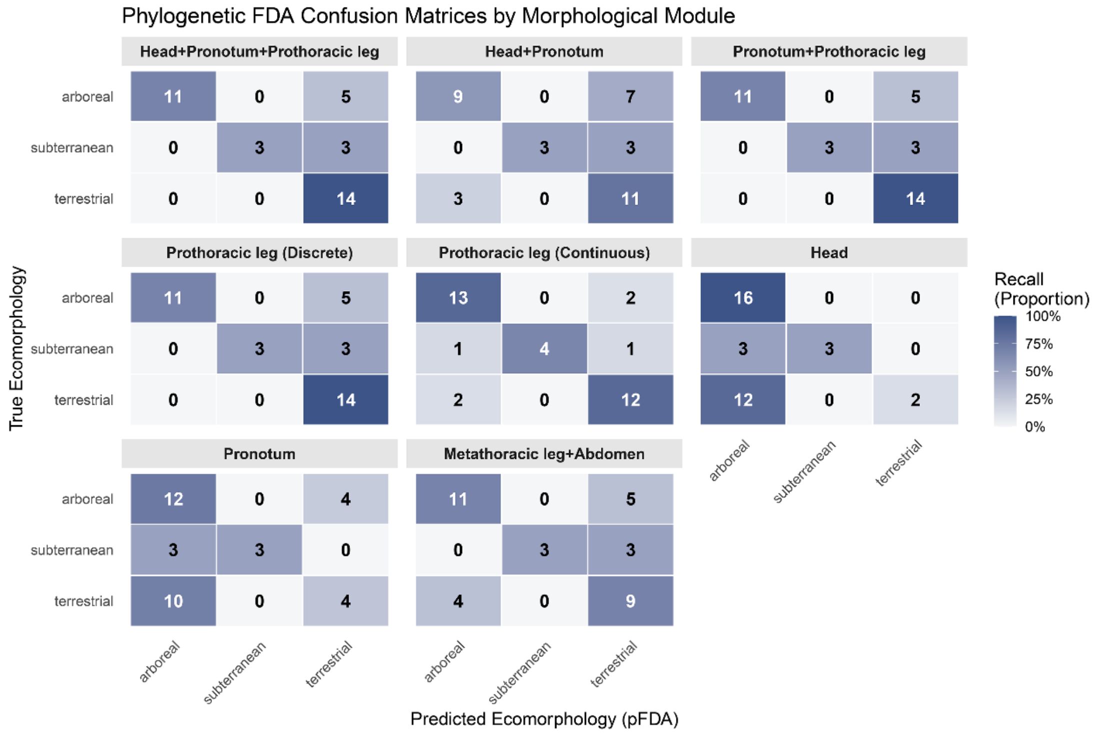


**Figure S3. Confusion matrices of phylogenetic flexible discriminant analysis (pFDA) models across eight morphological modules.** Ecomorphological classifications were evaluated using distinct morphological partition strategies: Head+Pronotum+Prothoracic leg, Head+Pronotum, Prothoracic leg (Discrete characters), Pronotum+Prothoracic leg, Head, Pronotum, Prothoracic leg (Continuous measurements), and Metathoracic leg+Abdomen. Rows represent true ecomorphological categories; columns denote pFDA predictions. Numerical labels indicate absolute species counts. Cell color gradients scale with the row-wise proportion (recall).

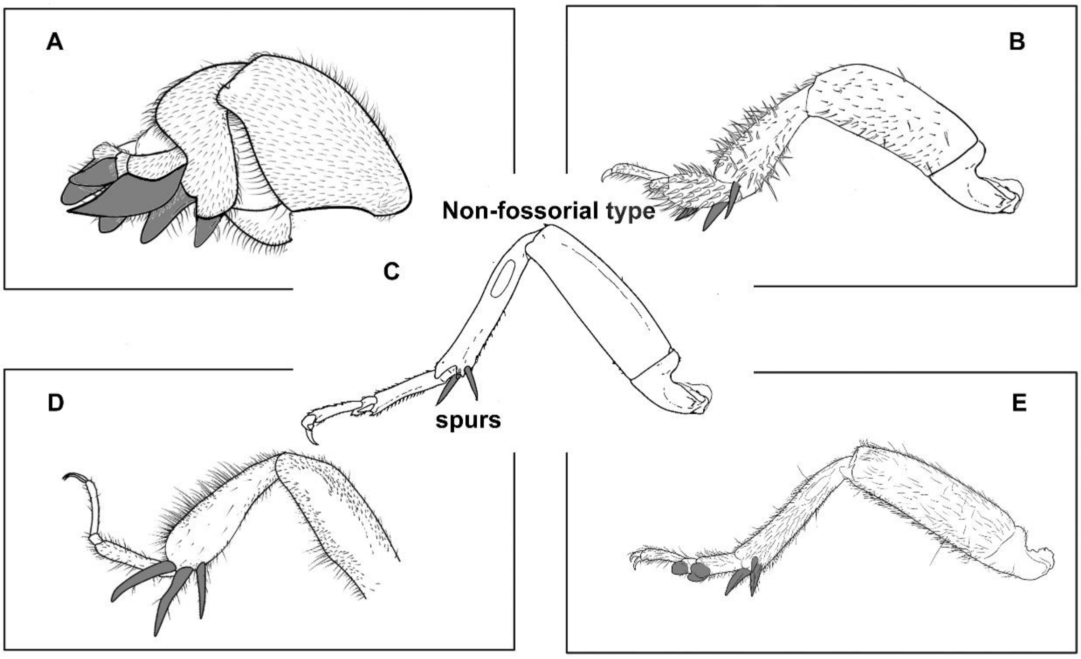


**Figure S4. Forelegs of five Gryllidea species, illustrating fossorial (A, B, D, E) and non-fossorial (C) types.** (A) *Gryllotalpa orientalis* Burmeister, 1838; (B) *Mellogryllus mutus* Cadena-Castañeda, Tavares & Fernandes, 2022, line drawing reconstructed based on morphological characters described and figured in Cadena-Castañeda et al. (2022)[S1]; (C) *Teleogryllus emma* (Ohmachi & Matsuura, 1951); (D) hypothetical generalized foreleg of Pseudogryllotalpidae, reconstructed primarily from *Petilus zhengi* Gu, Yuan & Ma, 2024[S2]; (E) *Gymnogryllus odonopetalus* Xie & Zheng, 2003.

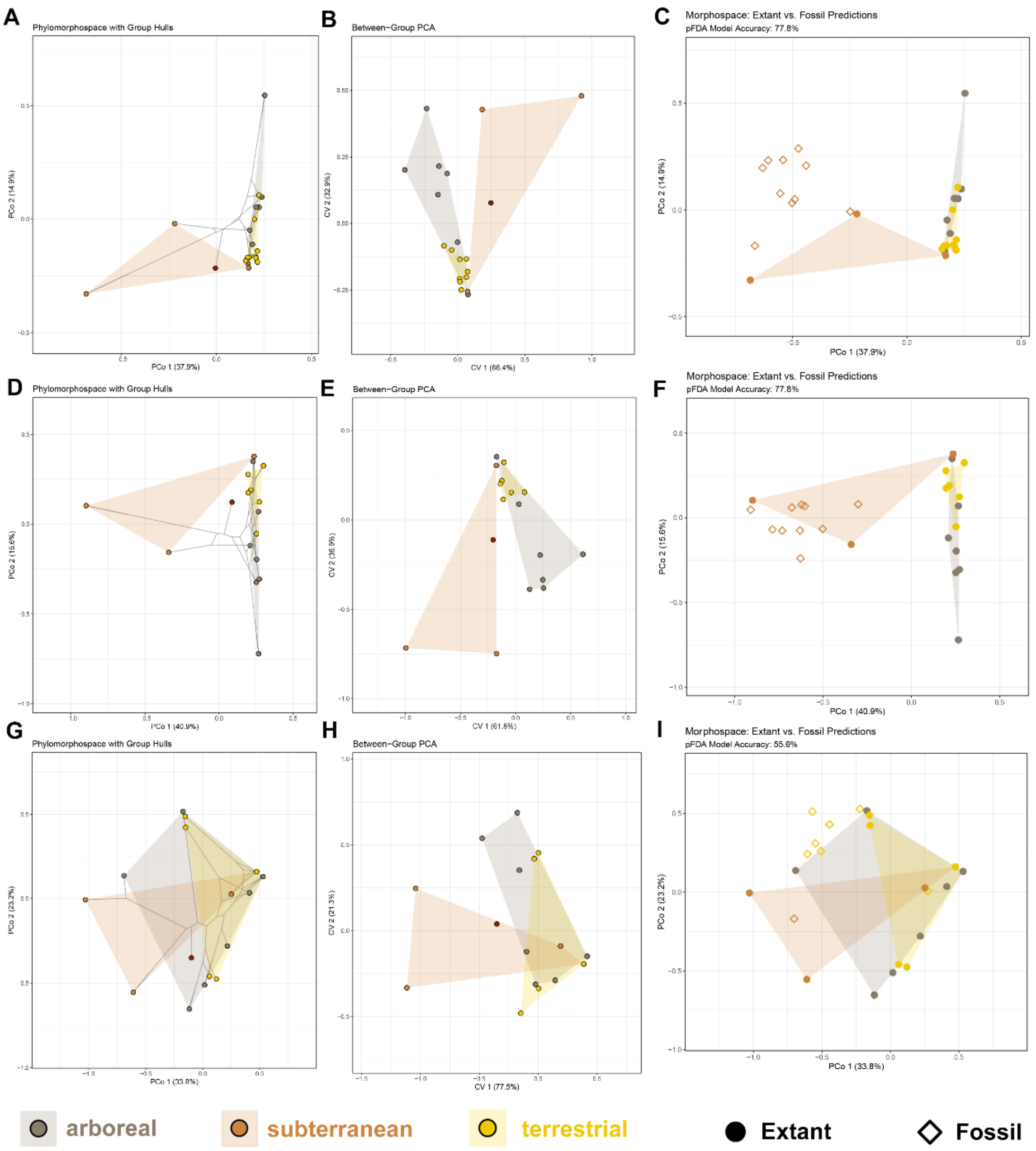


**Figure S5. Ecomorphological dynamics and niche predictions driven by the combined morphological partitions incorporating the prothoracic leg and the posterior module combining the metathoracic leg and abdomen partitions.** (A–C) Morphological disparity and ecological discrimination based on the combined head, pronotum, and prothoracic leg partition, illustrating the (A) phylomorphospace, (B) Between-Group PCA (bgPCA), and (C) phylogenetic Flexible Discriminant Analysis (pFDA). (D–F) Equivalent analyses, (D) phylomorphospace, (E) bgPCA, and (F) pFDA for the combined pronotum and prothoracic leg partition. (G–I) Equivalent analyses for the combined metathoracic leg and abdomen partition, illustrating the (G) phylomorphospace, (H) bgPCA, and (I) morphospace with pFDA-based predictions for fossil taxa.

### References


S1. O. J. Cadena-Castañeda, G. C. Tavares, J. A. M. Fernandes, Studies on Neotropical crickets: *Mellogryllus mutus* n. gen. et n. sp., an intriguing new genus and species of cricket of the Miogryllae Group (Orthoptera: Gryllidae: Gryllinae: Gryllini: Brachytrupina) from the Brazilian Atlantic Forest.

*Zootaxa* 5125, 408–420 (2022).
S2. J.-J. Gu, W. Yuan, L.-B. Ma, A. Nel, Z.-Q. Xu, N. Wang, C. Jiang, D. Ren, Y. Yue, More than a name: Mid-Cretaceous amber fossils link crickets and mole crickets (Orthoptera, Ensifera). *Syst. Entomol.* 49, 412–428 (2024).
